# Growth defect of domain III glycoprotein B mutants of human cytomegalovirus reverted by compensatory mutations co-localizing in post-fusion conformation

**DOI:** 10.1101/2023.03.28.534662

**Authors:** Mollik Madlen, Eisler Lukas, Külekci Büsra, Puchhammer-Stöckl Elisabeth, Görzer Irene

## Abstract

Cell entry is a crucial step for a virus to infect a host cell. Human cytomegalovirus (HCMV) utilizes the glycoprotein B (gB) to fuse the viral and host cell membrane upon receptor binding of gH/gL-containing complexes. Fusion is mediated by major conformational changes of gB from a metastable pre-fusion to a stable post-fusion whereby the central trimeric coiled-coils, formed by domain (D) III α helices, remain structurally nearly unchanged. To better understand the role of the stable core, we individually introduced three potentially helix-breaking and one disulfide bond-breaking mutation in the DIII α3 to alter the gB stability, and studied different aspects of the viral behavior upon long-term culturing. Two of the three helix-breaking mutations were lethal for the virus in either fibroblasts or epithelial cells and the third substitution led from mild to severe effects on viral replication and infection efficiency. gB_Y494P and gB_I495P suggest that the pre-fusion conformation was stabilized and the fusion process inhibited, gB_G493P on the other hand displayed a delayed replication increase and spread, more pronounced in epithelial cells, hinting at an impaired fusion. Interestingely, the disulfide bond-breaker mutation, gB_C507S, performed strikingly different in the two cell types – lethal in epithelial cells and an atypical phenotype in fibroblasts, respectively. Replication curve analyses paired with the infection efficiency and the spread morphology suggest a dysregulated fusion process which could be reverted by second-site mutations mapping predominantly to gB DV. This underlines the functional importance of a stable core for a well-regulated DV rearrangement during fusion.

**Importance:** Human cytomegalovirus (HCMV) can establish a lifelong infection. In most people, the infection follows an asymptomatic course, however it is a major cause of morbidity and mortality in immunocompromised patients or neonates. HCMV has a very broad cell tropism, ranging from fibroblasts to epi- and endothelial cells. It uses different entry pathways utilizing the core fusion machinery consisting of glycoprotein complexes gH/gL and gB. The fusion protein gB undergoes severe rearrangements from a metastable pre-fusion to a stable post-fusion. Here, we were able to characterize the viral behavior after the introduction of four single point mutations in gBs central core. These led to various cell type-specific atypical phenotypes and the emergence of compensatory mutations, demonstrating an important interaction between domains III and V. We provide a new basis for the delevopment of recombinant stable pre-fusion gB which can further serve as a tool for the drug and vaccine development.

## Introduction

Glycoprotein B (gB) is the highly conserved class III fusogen of herpesviruses [1]. The three dimensional conformations of pre- and post-fusion gB structures of different herpesviruses are well conserved [2] despite limited sequence similarity. Human cytomegalovirus (HCMV) UL55 encodes the 907 amino acid (aa) gB polypeptide (numbering according to TB40/E). Processing by proteolytic cleavage at the furin site [3, 4] results in an amino-(∼116 kDa) and carboxy-terminal (∼58 kDa) fragment, held together by two intramolecular disulfide bonds between cysteine residues C94 and C551 in domain (D)IV, as well as C111 and C507 in DIII [3]. gB is a membrane-bound homotrimer whose trimeric coiled-coils are formed by the α3 helices of DIII, centered on the three-fold axis of the ectodomain generating the trimerization contacts of the protomers [1, 5-7]. While comparison of gB pre- and post-fusion structures reveal major structural rearrangements during the transition process, the central core of the protein seems to remain nearly unchanged [6] indicating the importance of this stabilisation for fusion itself.

Herpesvirus gB is non-autonomously fusogenic [8, 9]. Instead, it requires activation by gH/gL-based glycoprotein complexes [10] which in HCMV is further linked to the accessory proteins gO (trimer) or UL128, UL130, and UL131A (pentamer) [11]. A current model of HCMV cell entry suggests that either the trimer, the pentamer, or both interact with pre-fusion gB [6, 12]. Binding of a gH/gL complex to the host cell receptor brings the viral and cellular membrane to close proximity for the gB fusion loop to insert into the membrane. Here, gB undergoes a major conformational change from a metastable pre-fusion to a stable post-fusion conformation resulting in the merging of the two membranes and the release of the viral content into the cell [1, 6]. Fusion at the plasma membrane of fibroblasts is pH-independent [13] and in the endosome of epithelial, endothelial, and myeloid cells pH-dependent [14]. After entry, a strict cell-to-cell spread was observed in clinical isolate culturing [15] which also extends to the dissemination of the virus in the human host [16, 17]. Cell culture-adapted strains, however, have at least partially lost this spread mode due to genomic alterations which mostly occur in the accessory proteins of the pentamer or in the RL13 locus [18, 19]. gB and gH/gL complexes are required for both, the cell-free and the cell-to-cell spread as for example small inhibitors directed towards gO of the trimer [20, 21] or towards gB [22] are able to inhibit both modes of spread. The pentamer can alternatively be used for cell-associated spread on fibroblasts and is sufficient for epithelial cell spread even without the presence of the trimer [23-26]. In clinical HCMV isolates, however, both gH/gL complexes are always present on virions [27].

Among the seven glycoproteins of the core fusion machinery gB, gH, and gO, are the most diverse expressing 5, 2, and 8 distinct genotypes, respectively, while gL, UL128, UL130, and UL131A show limited sequence diversity [28]. Those genotype sequences are widely distributed in the human population as they can be found in numerous geographically and clinically distinct populations [28-31]. Up to date, no association of a specific genotype with a higher disease severity has been found. In clinical samples different genotype combinations of the core fusion complex exist [32, 33] but not all theoretically possible combinations may occur due to a strong linkage disequilibrium between gH and gO genotypes [34-36].

On the single gene level, mounting evidence states that gB-mediated fusion can be differentially modulated by variants of the individual core fusion components. As recently reported a unique variant of gB, 275Y of AD196 strain and 585G of VR1814 strain, contributes to increased fusogenicity due to inherently hyperfusogenic gB or altered interaction with a gH/gL complex [37]. Furthermore, the gB genotype variability in the furin cleavage site differentially influences the cleavage efficiency, and probably the gB fusion capacity [4]. gH polymorphism contributes to the generation of gH genotype-specific antibodies depending on the underlying HCMV strain [38] and gO polymorphism alters the neutralizing capacity of gH and gH/gL-specific monoclonal antibodies [36, 39]. Additionally, gO polymorphism may influence the trimer to pentamer ratio [40], an important determinant for cell type-specific infection efficiency and the mode of spread [41], and may also affect the fusion process. In the present study, we focused on the large α3 helix of DIII forming the stable central core of gB. We selected four aa residues, G493, Y494, I495, and C507 for single point mutagenesis in the background of TB40-BAC4-derived strains with three different gO genotypic sequences, GT1c, GT3 and GT1c/3 [42]. Helix-breaking mutations to proline (G493P, Y494P, I495P) may contribute to a stabilization of gB in the pre-fusion conformation similarly as recently shown for HSV-1 [43]. The disulfide bond-breaking mutation (C507S) may affect the gB protomer coherence upon furin cleavage important for the structural stability [3] of the homotrimer. Long-term culturing of BAC-derived gB mutants demonstrated mutation- and cell type-dependent growth alterations, however with no prominent influence of the gO genotypic form. Second-site mutations in gB emerged exclusively in fibroblast-derived gB_C507S mutants, predominantly located in DV, and were associated with a rescue of the infection capability on both cell types.

## Results

### 1. gB mutants reveal mutation and cell type-dependent growth impairment

The four point mutations in the α3 helix of gB DIII were individually introduced into the TB40-BAC4-luc strain (Figure 1, Table 1). Two independent transfections and their subsequent analyses were performed in parallel under the same cell culture conditions in both, human foreskin fibroblasts (HFFs) and epithelial cells (ARPE-19), with gB mutants harbouring the gO background GT1c and GT1c/3. In an additional experiment, gB mutants with gO_GT3_ background were transfected into HFFs. Passaging was performed after 9 and 22 days post transfection (dpt) in HFFs, and after 9, 22 and 63 dpt in ARPE-19 cells. Cell-associated and/or cell-free viral DNA was measured over a time period of up to 121 dpt until a cytopathic effect (CPE) ≥ 80% was seen in HFFs and ≥ 50% in ARPE-19 cells (Supplementary Table 1).

**Table 1.**
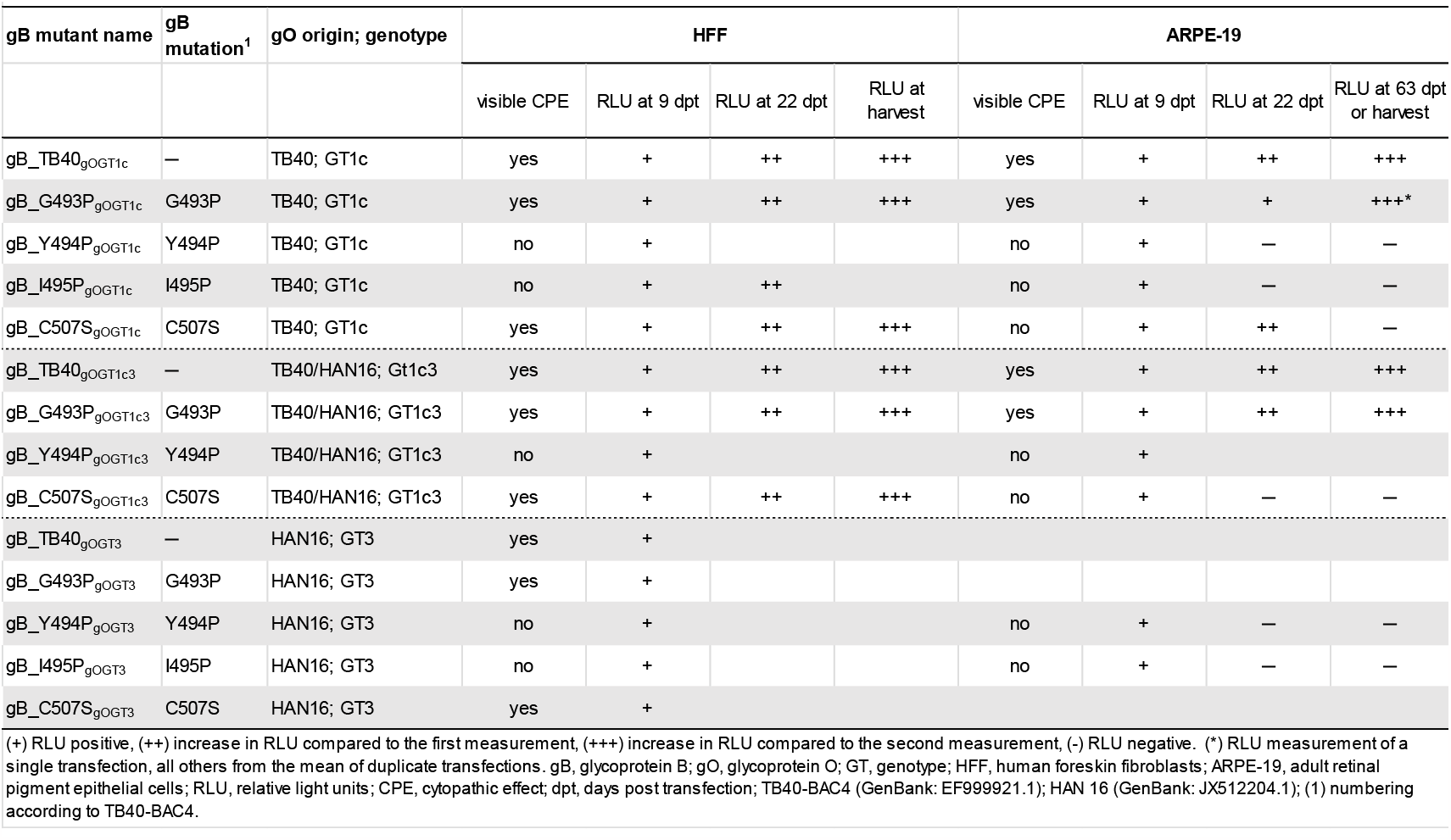
Transfection efficiency of parental and gB mutated viral genomes based on TB40-BAC4-luc into fibroblasts and epithelial cells.

| gB mutant name | gB mutation<sup>1</sup> | gO origin; genotype | HFF |  |  |  | ARPE-19 |  |  |  |
| --- | --- | --- | --- | --- | --- | --- | --- | --- | --- | --- |
|  |  |  | visible CPE | RLU at 9 dpt | RLU at 22 dpt | RLU at harvest | visible CPE | RLU at 9 dpt | RLU at 22 dpt | RLU at 63 dpt or harvest |
| gB_TB40<sub>gOGT1c</sub> | — | TB40; GT1c | yes | + | ++ | +++ | yes | + | ++ | +++ |
| gB_G493P<sub>gOGT1c</sub> | G493P | TB40; GT1c | yes | + | ++ | +++ | yes | + | + | +++* |
| gB_Y494P<sub>gOGT1c</sub> | Y494P | TB40; GT1c | no | + |  |  | no | + | — | — |
| gB_I495P<sub>gOGT1c</sub> | I495P | TB40; GT1c | no | + | ++ |  | no | + | — | — |
| gB_C507S<sub>gOGT1c</sub> | C507S | TB40; GT1c | yes | + | ++ | +++ | no | + | ++ | — |
| gB_TB40<sub>gOGT1c3</sub> | — | TB40/HAN16; GT1c3 | yes | + | ++ | +++ | yes | + | ++ | +++ |
| gB_G493P<sub>gOGT1c3</sub> | G493P | TB40/HAN16; GT1c3 | yes | + | ++ | +++ | yes | + | ++ | +++ |
| gB_Y494P<sub>gOGT1c3</sub> | Y494P | TB40/HAN16; GT1c3 | no | + |  |  | no | + |  |  |
| gB_C507S<sub>gOGT1c3</sub> | C507S | TB40/HAN16; GT1c3 | yes | + | ++ | +++ | no | + | — | — |
| gB_TB40<sub>gOGT3</sub> | — | HAN16; GT3 | yes | + |  |  |  |  |  |  |
| gB_G493P<sub>gOGT3</sub> | G493P | HAN16; GT3 | yes | + |  |  |  |  |  |  |
| gB_Y494P<sub>gOGT3</sub> | Y494P | HAN16; GT3 | no | + |  |  | no | + | — | — |
| gB_I495P<sub>gOGT3</sub> | I495P | HAN16; GT3 | no | + |  |  | no | + | — | — |
| gB_C507S<sub>gOGT3</sub> | C507S | HAN16; GT3 | yes | + |  |  |  |  |  |  |

(+) RLU positive, (++) increase in RLU compared to the first measurement, (+++) increase in RLU compared to the second measurement, (-) RLU negative. (\*) RLU measurement of a single transfection, all others from the mean of duplicate transfections. gB, glycoprotein B; gO, glycoprotein O; GT, genotype; HFF, human foreskin fibroblasts; ARPE-19, adult retinal pigment epithelial cells; RLU, relative light units; CPE, cytopathic effect; dpt, days post transfection; TB40-BAC4 (GenBank: EF999921.1); HAN 16 (GenBank: JX512204.1); (1) numbering according to TB40-BAC4.

**Figure 1.**
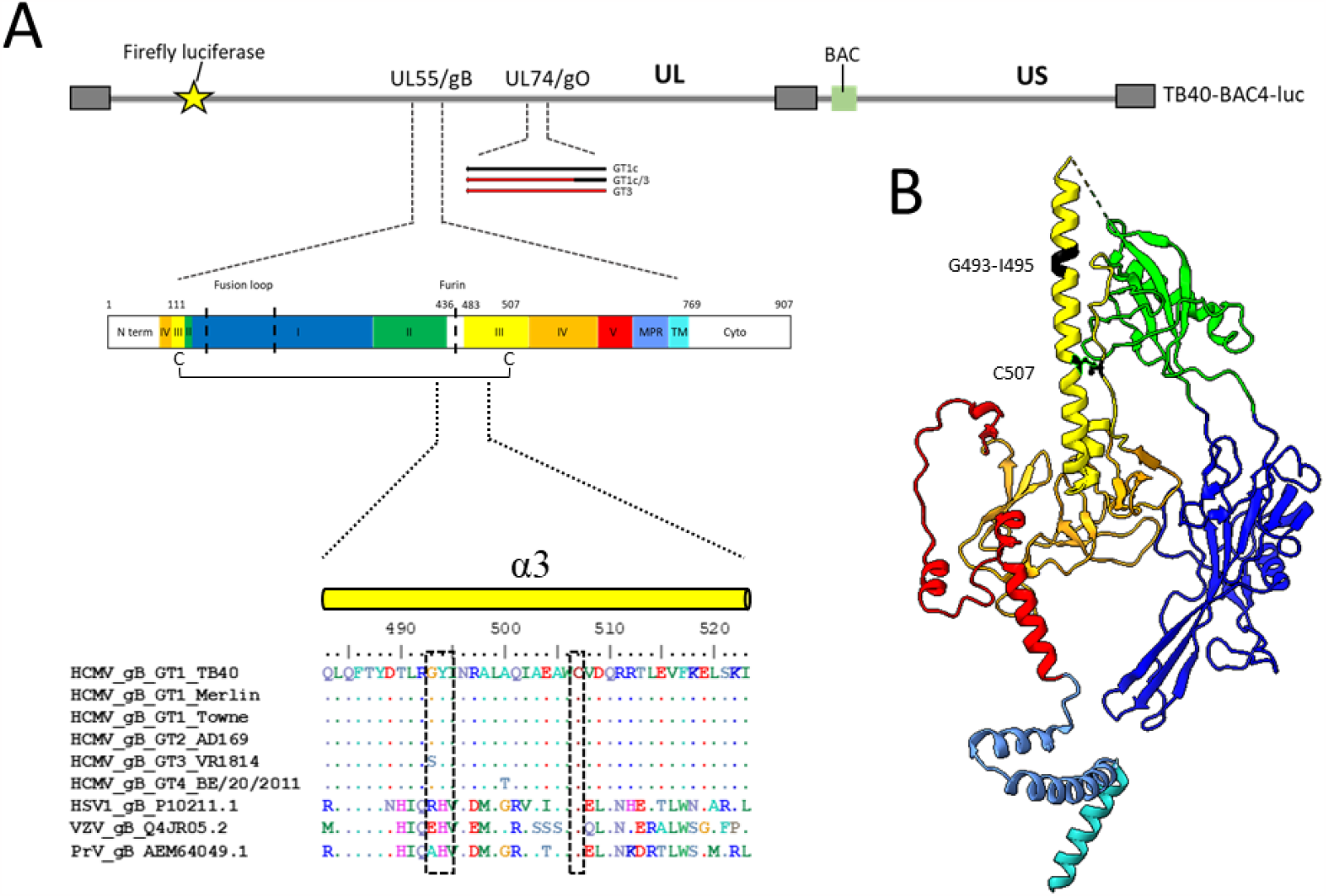
Schematic of TB40-BAC4-luc DNA and protomer gB pre-fusion structure. (A) Illustration of TB40-BAC4-luc DNA with main genome characteristics used for the transfection and reconstitution in fibroblasts and epithelial cells. Diagram of gB encoded by UL55 displaying the C111-C507 disulfide bond, fusion loops and furin cleavage site are marked with a dotted black line and the five gB domains (D) in color with the enlarged sequence alignment of the α3 helix with representative sequences of HCMV, HSV1, VZV, and PrV as indicated. The location of the mutations is depicted by dotted black boxes in the alignment. (B) Protomer structure (PDB: 5CXF). Target sites G493-I495 and C507, forming a disulfide bond with C111, are marked in black. gB domains are coloured as follows: DI (dark blue), DII (green), DIII (yellow), DIV (orange), DV (red), the membrane proximal region (MPR, blue) and the transmembrane domain (TM, turquoise) created with UCSF ChimeraX [61]. HCMV, human cytomegalovirus; GT, genotype; HSV1, herpes simplex virus 1; VZV, varizella zoster virus; PrV, pseudorabies virus; N-term, N-terminal region; MPR, membrane proximal region; TM, transmembrane domain; cyto, cytoplasmic tail.

First, the replication curves (as assessed by cell-associated and cell-free viral DNA) of gB_ G493P_gOGT1c_ and gB_ G493P_gOGT1c/3_ displayed a similarly slight delayed increase in the viral DNA load in the cellular and cell-free fraction of HFFs when compared to the parental strain (Figure 2A). No difference between the two clones of each mutant was seen. gB_ G493P_gOGT3_ showed a more severe delay in replication kinetics but displayed similarly high viral DNA loads as the parental strain at the time of harvest (Supplementary Figure 1). In ARPE-19 cells, gB_G493P_gOGT1c_ and gB_G493P_gOGT1c/3_ showed a severe replication delay (Figure 2B). The cell-associated viral load remained almost at the same level from 22 to 63 dpt and then increased 6-to 54-fold until harvest (between 108 to 121 dpt). Cell-free viral DNA started to steadily increase after 77 dpt. Interestingly, gB_G493P_gOGT1c/3_ clone 2 displayed the strongest viral load increase (2,5 log_10_ copies/ml) compared to gB_G493P_gOGT1c/3_ clone 1 (1,5 log_10_ copies/ml) and gB_G493P_gOGT1c_ clone 1 (1,9 log_10_ copies/ml). Notably, the replication delay was more pronounced in gB_G493P mutants carrying gO_GT1c_ compared to gO_GT1c/3_, and gB_G493P_gOGT1c_ clone 2 failed to establish long-term replication (Figure 2B).

**Figure 2.**
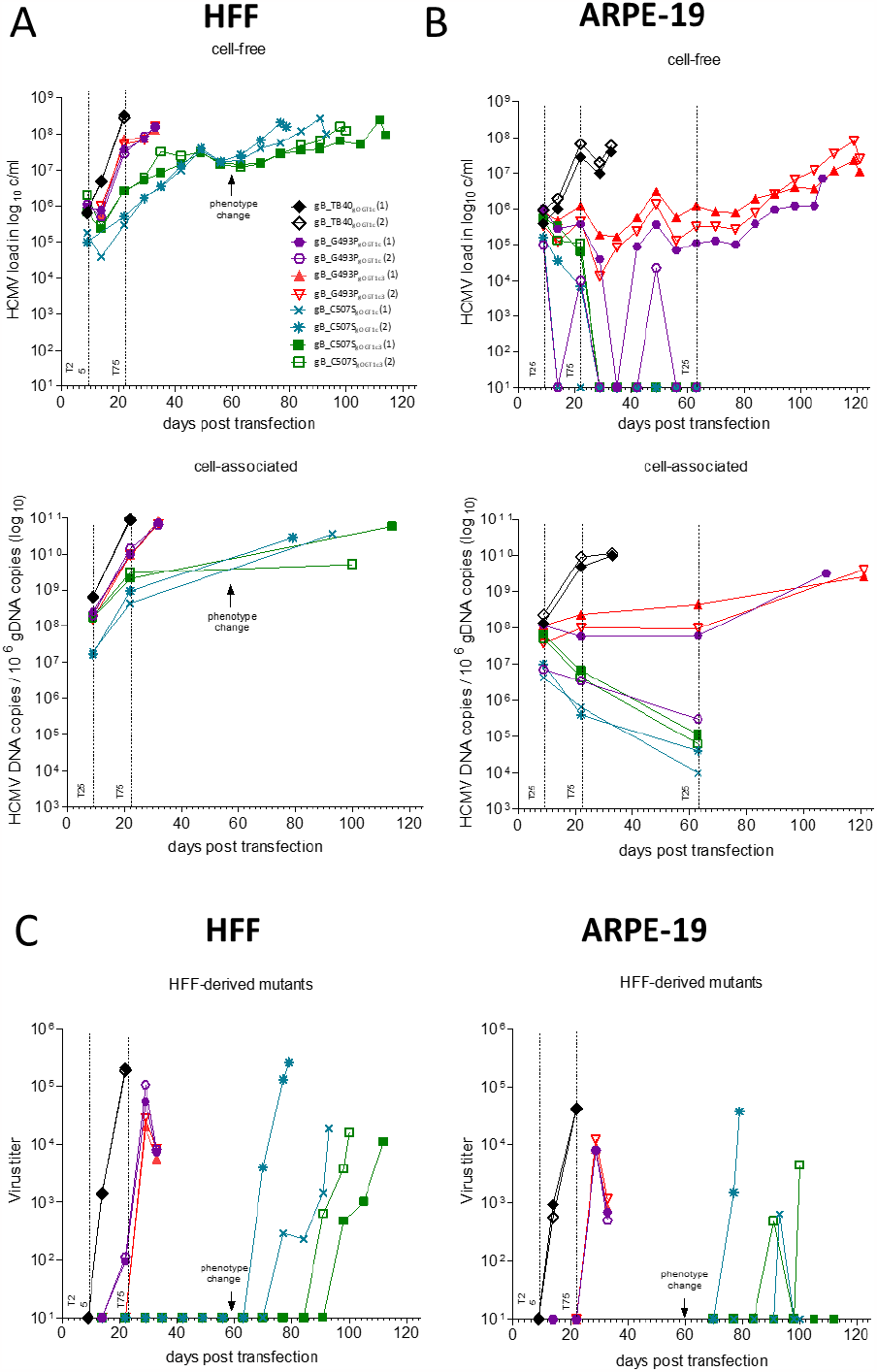
Viral replication and infectivity of gB mutants. A, B) Human foreskin fibroblasts (HFF) or epithelial cells (ARPE-19) were individually transfected in duplicates with the respective TB40-BAC-luc DNA and long-term cultured. During reconstitution, cell-free samples were taken from the supernatant once a week during medium change and cell-associated samples were taken during passaging as indicated. A) and B) top panel - viral DNA released into the supernatant of transfected HFF (A) or ARPE-19 (B) cells with the parental gB_TB40gOGT1c in black, gB_G493P mutants in red and purple, and gB_C507S mutants in green and petrol, each in gOGT1c and gOGT1c3 background, respectively. A) and B) Bottom panel - viral DNA load in cell-associated samples of transfected HFF (A) or ARPE-19 (B) cells normalized to the genomic DNA. C) Subsequent to the harvest of all gB clones, infection efficiency of cell-free viruses was tested in parallel through a luciferase assay. After infection, HFFs were incubated for 48 h and ARPE-19 cells for 72 h. The virus titer (mean of duplicates) was determined through the conversion of the RLU to MOI and further into PFU. Infection efficiency of HFF-derived parental gB_TB40gOGT1c, gB_G493P and gB_C507S mutants of all cell-free samples on HFFs (left). Positive samples on HFFs were then tested on ARPE-19 cells (right). HCMV, human cytomegalovirus; HFF, human foreskin fibroblast; ARPE-19, retinal pigment epithelial cells; T25, transfer into a T25 flask; T75, transfer into a T75 flask; gDNA, genomic DNA; RLU, relative light units; MOI, multiplicity of infection; PFU, plaque forming units.

Second, the replication curves of gB_ C507S already displayed strongly delayed replication kinetics in HFFs (Figure 2A, Supplementary Figure 1) and even failed to replicate in ARPE-19 cells (Figure 2B) despite successful transfection (Table 1). In the cell-associated fraction of HFFs, the viral DNA of gB_C507S_gOGT1c_ and gB_C507S_gOGT1c/3_ readily increased from 9 to 22 dpt for a mean of 1,4 log_10_ copies/ml comparable to gB_G493P (mean of 1,7 log_10_ copies/ml) and slightly lower than the parental strains (mean of 2,2 log_10_ copies/ml) (Supplementary Figure 2). The accompanied release of viral DNA (from 14 to 22 dpt) was significantly lower in gB_C507S (mean of 1,1 log_10_ copies/ml) than in parental strains (mean of 2,5 log_10_ copies/ml). Moreover, after a steady release of viral DNA for around 40 dpt, a stagnation period for 14 to 21 days set in, followed by an increase in viral load with a peak at >10^8^ copies/ml at harvest (between 79 and 114 dpt for gB_C507S_gOGT1c_ and gB_C507S_gOGT1c/3_) (Figure 2A). gB_C507S_gOGT3_ displayed a prolonged stagnation period (∼ 30 days), thus was harvested after 70 dpt (Supplementary Figure 1). Reinfection and further culturing of cell-free viruses for another 30 to 47 days led to a steady release of viral DNA.

Third, gB_Y494P and gB_I495P with the different gO genotypical backgrounds were unable to establish any detectable increase in viral DNA in neither cell type despite the proof of successfully transfected BAC DNA and repeatedly performed transfections (Table 1). Hence, no further culturing experiments were performed.

In summary, replication curve analyses upon transfection revealed striking differences among the individual gB mutants and cell types but without an obvious influence of the corresponding gO genotypes.

### 2. Fibroblast-derived gB_C507S are infectious for fibroblasts and epithelial cells after phenotype change

Due to the known producer cell effect, the capability of HFF-or ARPE-19 cell-derived virions to efficiently infect both cell types [44, 45], and in combination with the unknown effects of our introduced point mutations on the production of functional virions, we next determined the infection efficiency. For this, we performed an infectivity assay of our HFF- and ARPE-19 cell-released viruses (gB_G493P and gB_C507S) on both cell types (Figure 2C, Supplementary Figure 3A). All fibroblast-derived gB_G493P were able to infect HFFs and ARPE-19 cells. All fibroblast-derived gB_C507S gained the ability after the phenotype change to not only infect HFF but also ARPE-19 cells (except gB_C507S_gOGT1c/3_ clone 1) (Figure 2C) despite the lack of replication of gB_C507S upon transfection (Figure 2B). Initial detection of infectious virus was earliest for the parental strain (14 dpt), followed by gB_G493P (22 to 29 dpt), and was substantially delayed for gB_C507S (70 to 98 dpt in HFFs; 77 to 93 dpt in ARPE-19 cells). All the gB mutants had a lower infection capacity for ARPE-19 cells than for HFFs which is not significantly different from the ratio of the parental strain (Supplementary Figure 3B). This may explain why gB_C507S_gOGT1c/3_ with the lowest infection titer for HFFs did not lead to any detectable ARPE-19 cell infection. Infectious titer of ARPE-19 cell-released viruses was low for the parental strain and undetectable for the gB mutants in either cell type (Supplementary Figure 3A). All in all, fibroblast-derived gB_C507S attained infectiousness for both cell types while ARPE-19 cell-derived gB_G493P was incapable to establish an infection capacity.

### 3. Morphology of the cytopathic effect depends on the gB point mutation

The CPE of parental TB40-BAC4-luc virus is characterized by an even, supernatant-driven spread in HFFs while it is more restricted in ARPE-19 cells, as previously shown [44, 46]. Here, the CPE was visually inspected every 3 to 4 days by light microscopy over the whole time course. The morphologies markedly differed among the gB mutants and between the two cell types. In HFFs, all gB_G493P mutants except gB_G493P_gOGT3_ clone 1 showed a slightly delayed appearance of a parental strain-like CPE with an even spread of >90% CPE at harvest (Figure 3A, 3B; Supplementary Figure 4A, 4B). gB_G493P_gOGT3_ clone 1 presented a partially focal phenotype which appeared after 38 dpt for the first time and remained until harvest (Supplementary Figure 4B). In ARPE-19 cells, three of the four gB_G493P clones showed similarity to the parental strain, though the plaque formation was severely delayed while gB_G493P_gOGT1c/3_ clone 2 displayed one highly dense focus (Figure 3B). Due to disappearing CPE spots and a slow increase in the plaque sizes in ARPE-19 cells the two clones of gB_G493P_gOGT1c/3_ and gB_G493P_gOGT1c_ clone 1 (Figure 3B) were additionally passaged to reach >50% CPE at harvest. The dense foci of gB_G493P_gOGT1c/3_ clone 2 prevailed after passaging (Figure 3B). gB_G493P_gOGT1c_ clone 2 was not further cultivated as no viral DNA was detectable in the cells at 63 dpt (Figure 2B). In contrast to gB_G493P all clones of gB_C507S displayed a similarly aberrant spread pattern on HFFs (Figure 3C, Supplementary Figure 4C). After transfection a strictly focal CPE was formed and the foci appeared as highly dense accumulations of cells. From around 62 dpt onwards individual foci began to spread to the surrounding cells coinciding with the increase of released viral DNA (Figure 2A). This newly emerging CPE phenotype resembled the parental strain. At harvest, a combination of both led to a >90% CPE for gO_GT1c_ and gO_GT1c/3_ (Figure 3C) and to a >80% CPE for gO_GT3_ (Supplementary Figure 4C). No visible CPE was seen for gB_C507S in ARPE-19 cells consistent with their replication failure (Figure 2B). In overall, a variety of different spread phenotypes ranging from cell-associated to cell-to-cell pattern and a switch thereof could be observed during long-term culturing with no clear indication of a differential influence of the gO genotypes.

**Figure 3.**
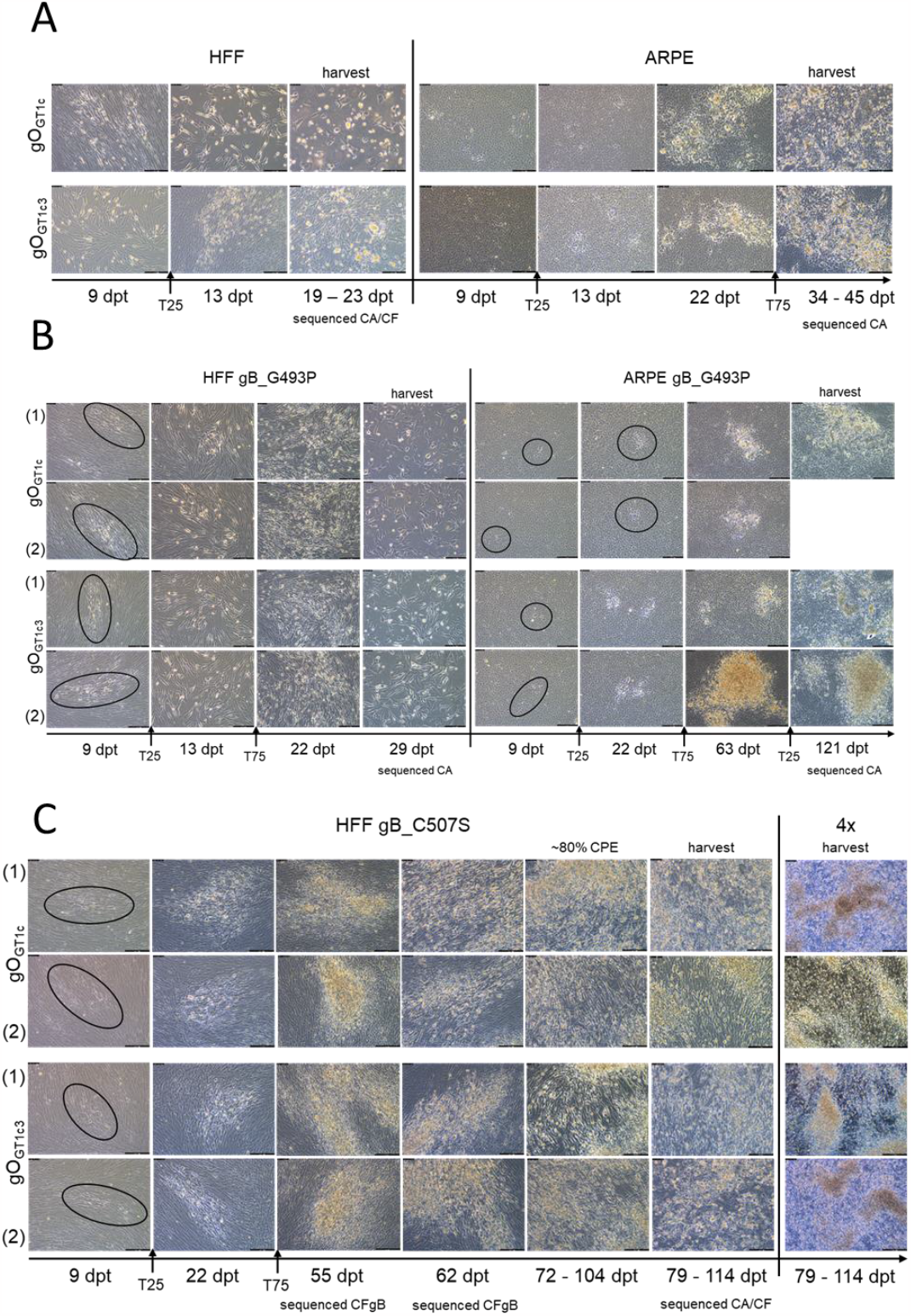
Spread morphology of gB mutants on fibroblasts and epithelial cells. Human foreskin fibroblasts (HFF) or epithelial cells (ARPE-19) were transfected with the respective TB40-BAC-luc DNA and long-term cultured. After transfection, pictures were taken every three to four days via light microscopy (Leica). Representative fields are shown at the indicated time points in 10X if not otherwise indicated. A) Parental strains gB_TB40gOGT1c (top) and gB_TB40gOGT1c3 (bottom) in HFF (left) and in ARPE-19 cells (right). Cell-free (CF) and cell-associated (CA) samples were sequenced at the time of harvest by whole genome sequencing (WGS). B) gB_G493P mutants in HFF (left) and ARPE-19 cells (right). WGS of CA samples at the time of harvest. C) gB_C507S mutants in HFFs. CA and CF samples were WGS sequenced at the time of harvest. CF samples at 55 and 62 days post-transfection (dpt) were enriched through long range PCR for gB (CFgB) prior to sequencing due to a low viral load. Black circles indicate cytopathic effects (CPE). HCMV, human cytomegalovirus; HFF, human foreskin fibroblast; ARPE-19, retinal pigment epithelial cells; T25, transfer into a T25 flask; T75, transfer into a T75 flask; (1), clone 1; (2), clone 2; dpt, days post transfection; CF, cell-free; CA, cell-associated; CPE, cytopathic effect; CFgB, cell-free gB.

### 4. gB_C507S rather than gB_G493P acquire second-site mutations in gB

The distinct CPE patterns prompted us to investigate the HCMV genome for the evolution of reversion and/or second-site mutations after long-term culturing post-transfection (parental strains, n=10; gB mutants, n=15) and after reinfection (gB_C507S_gOGT3_, n=2). Therefore, whole genome sequencing (WGS) was performed using the cell-associated and/or cell-free viral DNA collected at harvest. The HCMV-DNA load of >10^9^ log_10_ copies/ml was sufficiently covered without any further enrichment. All gB mutants maintained the introduced point mutations, while no mutations at all were observed for the parental strains in both cell types (Supplementary Table 1a). Eight out of 15 gB mutant clones acquired additional mutations at different genomic locations and at various frequencies, accompanied with a phenotype change (Supplementary Table 1b). gB_G493P_gOGT3_ clone 1 developed a single nucleotide polymorphism in an overlapping open reading frame in HFFs leading to UL76_L131I and UL77_S324Y substitutions at a frequency of ∼80%. gB_G493P_gOGT1c3_ clone 2 acquired a UL131A_T101S mutation in ARPE-19 cells with a frequency of almost 100%. No further genomic mutations were observed for other gB_G493P clones in both cell types and no second-site mutations developed in gB. This is in stark contrast to the fibroblast-derived gB_C507S, where all developed at least one additional mutation in gB (Table 2, Supplementary Table 1b). These substitutions were found at various frequencies (≤ 82%) and at different regions, yet particularly concentrated in DV (Figure 4A). Furthermore, gB_C507S_gOGT1c_ clone 2 and gB_C507S_gOGT1c3_ clone 2 each acquired genomic mutations in UL48_A1684V and UL148_D247A, respectively. Together, the emergence of the additional mutations seems to be associated with the type of initially introduced mutation and with the appearance of a phenotype switch rather than with a certain gO genotype.

**Table 2.**
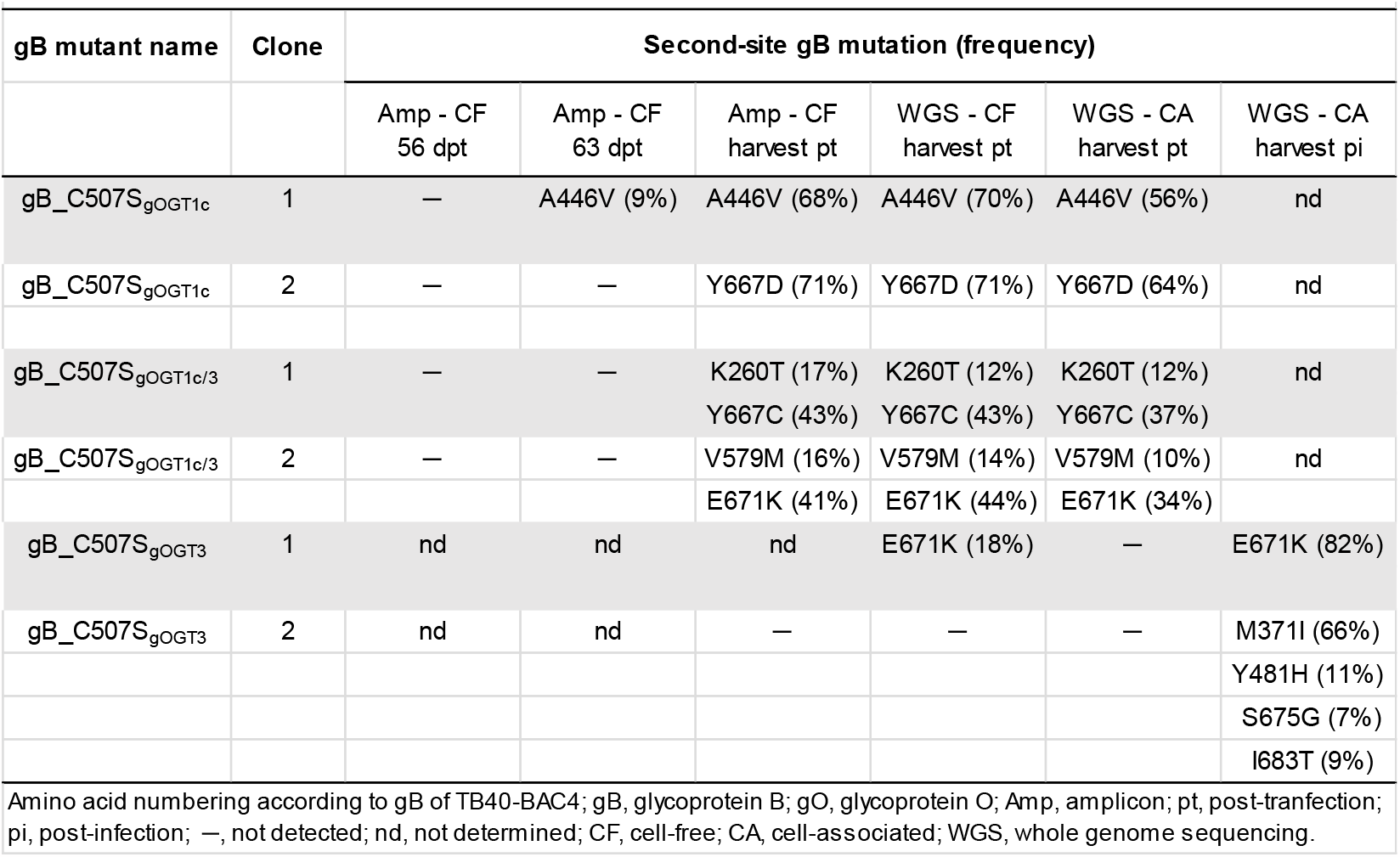
Appearance of second-site gB mutations in fibroblast-derived gB_C507S mutants detectable in cell-free and cell-associated fraction after transfection or reinfection.

**Figure 4.**
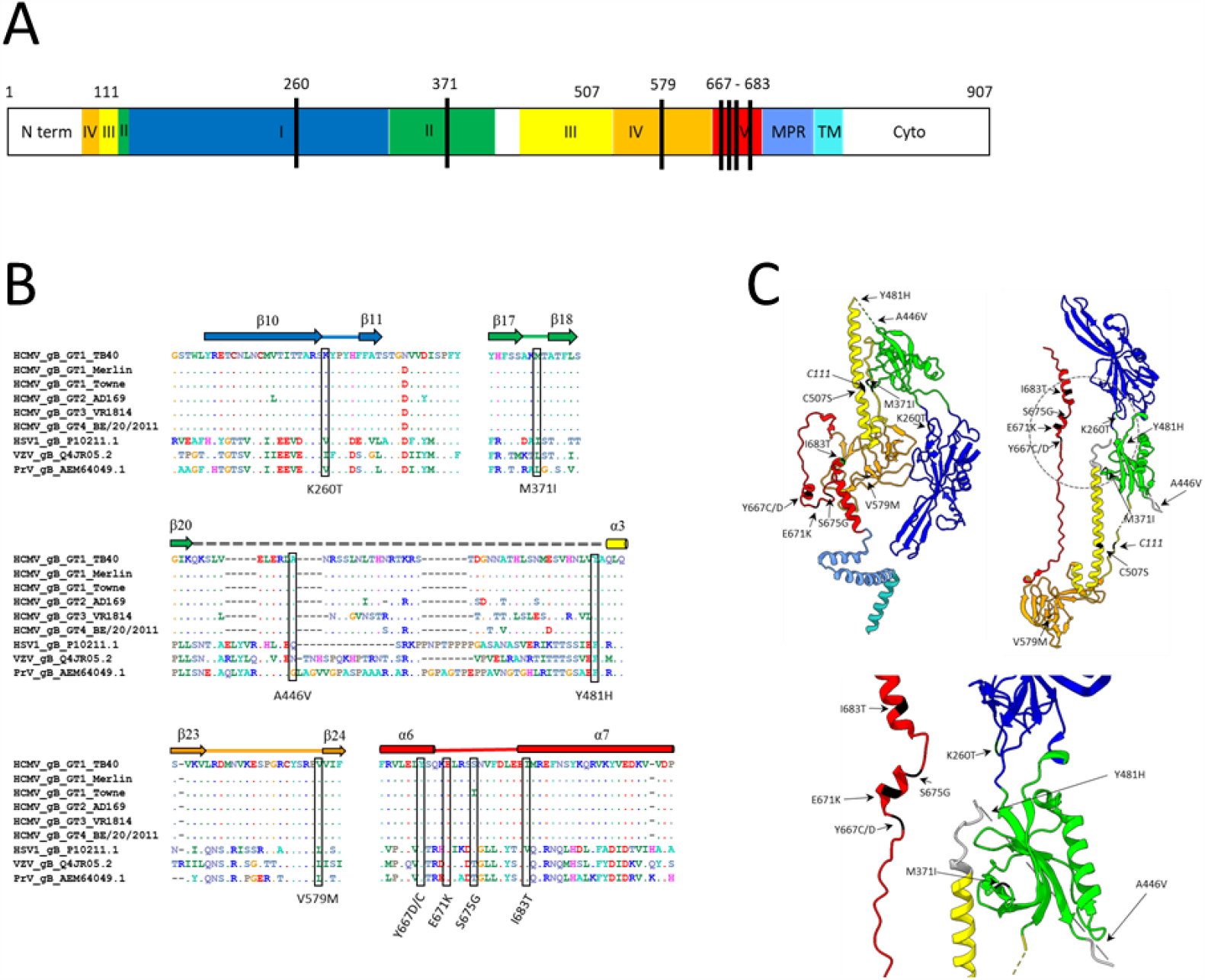
Location of gB second-site mutations in gB_C507S mutants after reconstitution. A) Schematic diagram of linear gB marked with acquired second-site mutations as black lines. B) Amino acid alignments of gB from different herpesviruses as indicated with the homologous sites of the second-site mutations highlighted with black boxes. Respective substitutions are shown below. C) Protomer structure of gB with domain (D) I (dark blue), DII (green), DIII (yellow), DIV (orange), DV (red), the membrane proximal region (MPR, blue) and the transmembrane domain (TM, turquoise) created with UCSF ChimeraX [61]. Target site C507S and C111 are marked in black. Acquired second-site mutations are highlighted in pre-fusion protomer (top left; PDB: 5CXF), post-fusion protomer (top right; PDB: 5C6T) and in a zoom of the encircled post-fusion region (bottom). HCMV, human cytomegalovirus; HSV1, herpes simplex virus 1; VZV, varizella zoster virus; PrV, pseudorabies virus; N-term, N-terminal region; MPR, membrane proximal region; TM, transmembrane domain; cyto, cytoplasmic tail.

### 5. Detection of second-site gB mutations briefly after the phenotype change

The observed reversion of the atypical to parental strain-like spread pattern in combination with the additional mutations in gB of fibroblast-derived gB_C507S at harvest led to the question if these were already detectable around the appearance of CPE change. Collected supernatants prior to (56 dpt) and shortly after the CPE change (63 dpt) were chosen from gB_C507S_gOGT1c_ and gB_C507S_gOGT1c/3_ and subjected to in-depth sequencing (Figure 3C). Due to low viral loads of ∼10^7^ copies/ml, the gB region was enriched by long-range (LR) amplicon prior to sequencing as previously described [35]. No second-site gB mutations were detectable before the morphological change (Table 2, Supplementary Table 1c), while in gB_C507S_gOGT1c_ clone 1 A446V appeared briefly afterwards at a frequency of 9%. This strongly suggests that the evolution of a second-site gB mutation accounts for the phenotype change of fibroblast-derived gB_C507S. In addition, cell-free and cell-associated virions carried the same mutational pattern with the same ratio of major and minor substitutions in WGS as well as amplicon sequencing.

## Discussion

Using targeted mutagenesis on BAC-derived viruses we demonstrate that single point mutations in the central α helix of DIII HCMV gB cause severely distinctive growth defects in fibroblasts and epithelial cells, reinforcing the mechanistic differences of gB-mediated fusion between these two cell types [47]. In vitro evolution upon long-term culturing reveals compensatory gB mutations that are predominantly concentrated in DV corroberating a cooperative regulatory role of DIII and V during fusion.

The most exceptional phenotype was displayed by gB_C507S on fibroblasts. Each α3 of the three gB protomers form a coiled coil structure leading to a stable central structure of the homotrimer which stays nearly unchanged during the pre-to post-fusion transition [6]. Further, C507 forms a disulfide bond with C111 to maintain the association of the N- and C-terminal gB fragments upon furin cleavage [3, 5]. We hypothesize that the disruption of this disulfide bond may weaken the stability, thereby affecting the fusion process. Our data show that gB_C507S display tightly localized foci similar to a cell-to-cell spread in fibroblasts that contrasts the evenly distributed cell-free spread of the parental TB40-BAC4-luc strain [44, 46]. These findings hint at an alteration of the fusion process by C507S rather than a global disruption of the gB function. Despite this obvious functional dysregulation the virions can spread as reflected by an increase in the focal CPE and in the cell-associated viral DNA, yet the virion release is substantially reduced.

Intriguingly, the focal growth changed to a parental strain-like spread coinciding with the evolution of additional mutations in gB, which indicates an, at least partial, compensation of the gB fusion defect. Even more remarkable is the number of compensatory mutations mapping to gB DV (Figure 4B-C) attributing an important functional relationship between DIII and DV. Recent determination of the pre-fusion structure of HCMV gB shows that DV, by contrast to DIII, undergoes substantial rearrangement during the conformational change [1, 6]. Refolding and extension of DV is needed for the transition of the extended intermediate to the post-fusion conformation. DV drags the membrane proximal region and transmembrane domain close to the fusion loops in DI, bringing the two membranes together. In the post-fusion conformation, the three α helices of DV (α5, α6, α7) form an extensive trimerization interface with DIII and DI. Hence, it can be speculated that the newly evolved mutations in DV compensate for the engineered instabilization which might be particularly disadvantageous in the last steps of the fusion process. The crucial role of DV as fusion regulator was previously shown for other herpesviruses [2]. Point mutations in the arm region, for example, cause a reduced fusion ability for human herpes simplex virus 1 (HSV-1) [48], revertable by V705I and N709H in α7 [49]. Further, N735S in α7 of pseudorabies virus (PrV; *Suid alphaherpesvirus 1*) reverts an entry-deficient mutant back to hyperfusogenity [50]. The variant allele Q/R699, localized in α6 of varicella-zoster virus (VZV), of which Q699, dominant in the live varicella Oka vaccine and a determinant for its attenuation, showed significantly lower fusion activity than R699 [51]. Here, we identified various mutations localized within or in proximity of α6 and α7 with I683T homologous to V705I in HSV-1 (Figure 4 A-C) [49]. Together, these findings underline the key role of the proper rearrangement of DV in regulating the fusogenic ability of gB not only in α-but also in β-herpesviruses.

Upon phenotype change and acquirement of additional mutations in gB_C507S, the virus release increased and virions became infectious for both, fibroblasts and epithelial cells -with an epithelial to fibroblast ratio resembling the parental strain [44] – indicating a reversion of the mutant phenotype. However, the increase in release was not as fast as expected even after reinfection, which suggests that the additional mutations are unable to completely rescue the mutant phenotype. Moreover, our deep sequencing data revealed that none of the second site gB mutations reached near 100% dominance while the originally introduced C507S remained stable.

The finding that gB_C507S alone completely abolishes growth on epithelial cells in contrast to fibroblasts reflects the mechanistic differences in the fusion process. Virus entry into fibroblasts occurs under neutral pH at the plasma membrane and into epithelial cells under low pH at the endosomal membrane [13, 14]. We assume that exposure to low pH further alters the impaired HCMV gB structure, as shown for HSV-1 gB [52], leading to these drastic effects on epithelial cells. Although the furin cleavage site is highly conserved among herpesviruses very little is known about the gB maturation process in HCMV [4, 53]. We speculate that HCMV uses furin cleavage only during the endosomal entry and not for direct plasma membrane fusion. Here, gB_C507S would underline this hypothesis due to the contrasting growth phenotypes in both cell types. Another not mutually exclusive explanation relies on the cell type-specific requirements of the distinct gH/gL complexes for gB triggering [12, 54, 55]. Either trimer or pentamer can mediate virus spread on fibroblast while pentamer is sufficient on epithelial cells [23-26]. Hence, we hypothesize that gB_C507S affects the interaction with the pentamer stronger than with the trimer. Consequently, an in-depth analysis can provide better understanding of the gB-mediated fusion triggers in both, cell types and complex interactions.

The polymorphic subunits of the core fusion machinery, in particular gB, gH, and gO [28], may also influence the fusion process. Here, we compared gB mutant phenotypes in three gO genotypic backgrounds, gO_GT1c_, gO_GT3_, and the chimera gO_GT1c/3_ [42]. gO_GT1c/3_ virions carry a substantially lower content of trimer than gO_GT1c_ and gO_GT3_, while gO_GT3_ and gO_GT1c/3_ infect epithelial cells better. We assumed that distinct growth properties may differentially contribute to effects of gB mutations. Here, we were unable to see an obvious gO-dependent effect on mutant phenotypes, however we solely used one genotype for gB and gH. To further clarify whether or to which extent various combinations, as seen in HCMV clinical strains [33], influence gB-mediated fusion need to be tested. Inspection of all available HCMV gB sequences shows that Y494 and I495 are highly conserved while around 30% carry a serine instead of a glycine at site 493 (gB genotype 3). This polymorphism may explain why a substitution to proline has a less severe effect on the gB function. gB_G493Pallows growth on both cell types whereas gB_Y494P and gB_I495P completely abolish virus growth pointing to a complete loss of gB function. Y494 maps to the homologous residue H516 in HSV-1 where a proline substitution leads to functional arrest in the pre-fusion conformation [43]. If the loss of function in HCMV is due to a pre-fusion arrest or due to effects on the expression, folding, processing or transport awaits further investigations. In contrast, gB_G493P grew almost parental strain-like on fibroblasts and with a severe growth retardation on epithelial cells. We assume that also a low pH and/or differences in the gB processing and/or the interaction with the different gH/gL complexes further weakens the gB function or structure. gB_G493P lack any emergence of second-site gB mutations, hinting at an impaired fusion process contrasting to gB_C507S with a dysregulated fusion, especially visible on epithelial cells. Consequently, the G493P defect is unable to be rescued by mutations in gB.

Additional mutations at other genomic locations were sporadically observed in gB_C507S and gB_G493P on either cell type. Exceptionally, UL131A_T101S evolved with around 100% frequency in one gB_G493P clone in epithelial cells implying a growth advantage over the initial mutant. The associated formation of large dense foci may reflect a change in the mode of virus spread. UL131A, a highly conserved component of the pentamer [56, 57], plays a role in virus release [58, 59], and mutations thereof influence the mode of spread due to a change in the ratio of gH/gL complexes [18, 19]. From our data it appears that the emergence of T101S renders the viruses more efficient to cell-to-cell spread and to virus release which is well in line with the predicted functions of UL131A. In summary, we have introduced single point mutations situated in the DIII α3 helix in the HCMV gB and characterized the viral behavior in long-term evolutionary culturing. We underline the importance of DIII and DV during the transition process on which future studies could focus to better understand the cell entry and especially the mode of spread in different cell types. Finally, we identified a potential for the production of recombinant stable pre-fusion gB which could serve as a tool for the vaccine and drug development.

## Materials and Methods

### Cells

Human foreskin fibroblasts (HFFs) and human adult retinal pigment epithelial (ARPE-19; ATCC, Manassas, Virginia) cells were cultivated at 37°C, 5% CO_2_ in minimum essential medium Eagle (MEM, Sigma Life Science) supplemented with 10% Fetal Calf Serum (FCS, Capricorn Scientific) and 0.5% Neomycin (Sigma-Aldrich).

### Generation of gB mutants

Previously generated bacterial artificial chromosome (BAC) TB40-BAC4-luc DNA [44] carrying either gO genotype (GT) 1c, GT1c/3 or GT3 sequence [42] were used. Point mutations were individually introduced via en passant mutagenesis [60] into gB of the respective BAC DNAs using *Escherichia coli* (*E. coli*) GS1783 strain as previously described [46]. Briefly, the recombination cassette was amplified from pEP-Kan-S (kindly provided by N. Osterrieder) by PCR using mutagenesis primer (Supplementary Table 2). After electroporation, recombination-positive *E*.*coli* were subjected to kanamycin selection to remove the non-HCMV sequences within *E*.*coli* by cleavage at the I-Sce I site and a second red recombination. For BAC-DNA purification the NucleoBond™ Bac100 kit (Macherey-Nagel) was used following the manufacturer’s protocol. BAC-DNA pellet was dissolved in 400 μl deionized water, mixed by flicking the tube and incubated for 2 h at RT. The BAC-DNA was stored at 4°C for up to 2 weeks until subjected to whole genome sequencing (WGS) and reconstitution.

### Transfection, Reconstitution, and long-term culturing

HFF and ARPE-19 cells were seeded on a 6-well plate (1 – 1.5 × 10^5^ cells/ml) 24 h prior to the transfection. The following day, medium was removed and 2.7 ml of fresh MEM with 10% FCS, without antibiotics, were added. For transfection 2.5 μg BAC-DNA for all mutants except for gB_G493P_gOGT1c_ clone 1 (2.0 μg) and gB_G493P_gOGT1c_ clone 2 (3.0 μg) were used. The transfection mix further included 0.5 μg of CMV71 plasmid (kindly provided by Mark Stinski), 12 μl Viafect Transfection Reagent (Promega) and MEM without antibiotics and FCS to a final volume of 300 μl. After a 10 min incubation at RT the transfection mix was transferred to the cells and after 24 h, the medium was exchanged with fresh MEM with 0.5% Neomycin and 10% FCS. Cells were passaged into T25 flasks after 10 – 13 days and into T75 flasks after 23 – 26 days. ARPE-19 cells were additionally passaged into T25 flasks after 63 days. Medium change was performed up to twice a week. Two to four independent transfections were performed for each BAC mutant.

### Viral Growth During Reconstitution

Cytopathic effect (CPE) in cell culture monolayer was visualized using a 4X or 10X objective with the Leica MC170HD microscope with integrated camera and the Leica Application Suite X (LASX) program every 3 – 4 days until virus harvest. Cells and supernatant were harvested at a 95 – 100% CPE in HFFs and at around 50% CPE in ARPE-19 cells. At each medium change the supernatant was cleared by centrifugation for 20 min at 4000 rpm, 4°C. One aliquot (2μl) was immediately transferred to 2 ml lysisbuffer for DNA extraction using the bead-based NucliSens EasyMag extractor (BioMérieux) according to the manufacturer’s protocol, eluted in 50 μl of nuclease-free H_2_O and HCMV-DNA quantified. Residual 1ml aliquots were stored at –80°C before subjected to long-range PCR, sequencing and infectivity testing. During each passaging step and at harvest, a 500μl aliquot of the cell suspension was transferred to 2 ml lysisbuffer for DNA extraction, HCMV-DNA and genomic beta-globin DNA quantification. Another 300μl of the cell suspension were used for determination of relative light units (RLUs) through a luciferase assay. Extracted DNA from cell-associated samples were stored at -20°C before subjected to WGS.

### Luciferase assay

Positively transfected BAC DNA was confirmed by firefly luciferase assay. Equal parts of Steady-Glo (Promega) substrate were added to the cell suspension, incubated for 5 min at RT and luminescence measured with a Synergy HTX multimode reader (BioTek) in triplicates.

### HCMV DNA-, beta globin-specific qPCR and long-range PCR

For HCMV DNA quantification, a region within US17 gene and for quantification of cellular genomic DNA (gDNA), a region within the human beta-2-microglobulin gene was amplified as previously described [46]. Enrichment of the gB gene including up- and downstrean genomic regions was performed by long range PCR with the primer pairs, CMVF1_new, ACCAGATGCTGACGATAGC, and CMVR1.1, TCCAGAACGTGGCGCTTATT generating a 7956 bp amplicon as previously described [35].

### Next generation sequencing

Isolated BAC DNA, total DNA extracted from transfected cells and cell culture supernatant, and long range amplicons were used as input DNA for library preparation using the Nextera XT DNA Kit (Illumina) following the manufacturer’s protocol. In short, 2 ng input DNA was used to yield a 4 nM library, which was spiked with 15 pM PhiX (2.5%) as control. Samples were sequenced using the MiSeq™ sequencer (Illumina) with paired-end reads (2×150-250) on a MiSeq system using v2 sequencing reaction chemistry (Illumina). The resulting sequencing reads were analyzed in CLC Genomics 21.0 (Qiagen). The average coverage was calculated by the average length of mapped reads multiplied by the number of mapped reads and divided by the reference genome length of TB40-BAC4-luc.

### Infectivity Assay

For measuring the infection efficiency of the viral particles during the period of reconstitution, HFF and ARPE-19 cells were seeded in 96-well plates (1 – 1.5 ×10^5^ cells/ml). The next day, 100 μl cleared supernatants from all time points of medium change were added to the medium with one additional serial dilution of 1:2 to infect HFF and incubated for 48 h at 37°C and 5% CO_2_. ARPE-19 cells were infected with cleared supernatants which prior tested positive on HFF cells and incubated for 72 h at 37°C and 5% CO_2_. Then medium was removed, cells washed once with 1X PBS and lysed with 100 μl Glo Lysis Buffer (Promega) and incubated for 5 min at RT. Plates were stored at -20°C before adding 100 μl Steady Glo substrate (Promega) to perform a luciferase assay as described earlier. Results were shown in RLU. The viral titer was calculated with a previously tested correlation of RLU to multiplicity of infection (MOI). The following formulas were used to calculate plaque formation unit (PFU) per ml: MOI = RLU * correlation factor; PFU = MOI * number of seeded cells.

## Acknowledgments

We kindly acknowledge the support and technical assistance of Andreas Rohorzka, Michaela Binder, Barbara Dalmatiner, Birgit Gangl, Sylvia Malik and Gabriele Sigmund.

The authors declare no conflict of interest.

## Supplementary Material

**Supplementary Figure 1.**
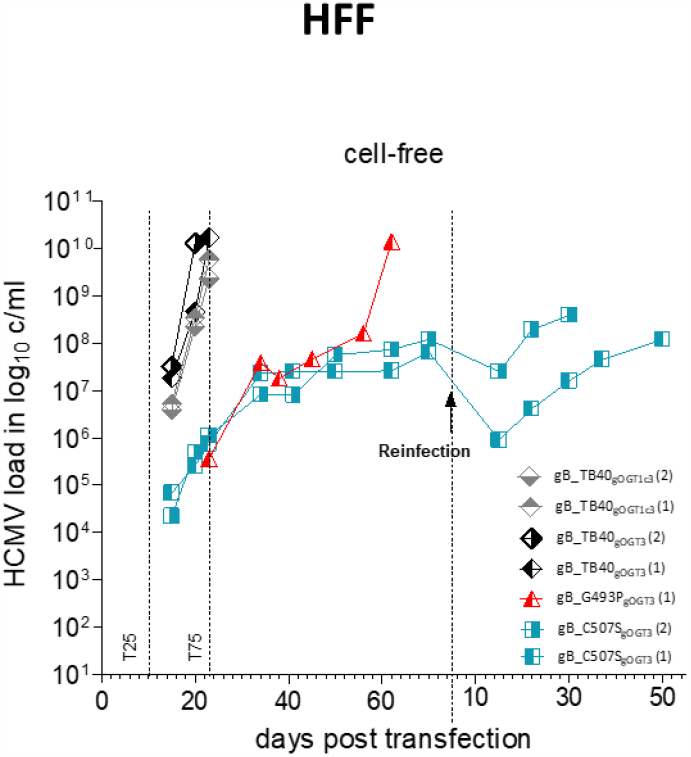
Viral replication curves during reconstitution and reinfection of cell-free samples. Fibroblasts (HFF) were transfected with the respective TB40-BAC-luc DNA and long-term cultured. During reconstitution, cell-free samples were taken from the supernatant once a week during medium change. Viral DNA released into the supernatant of the parental gB_TB40_gOGT3_ in black, gB_TB40_gOGT1c3_ in grey, gB_G493P mutant in red and gB_C507S mutants in petrol, each in gOGT3 background. HCMV, human cytomegalovirus; HFF, human foreskin fibroblast; GT, genotype; T25, transfer into a T25 flask; T75, transfer into a T75 flask.

**Supplementary Figure 2.**
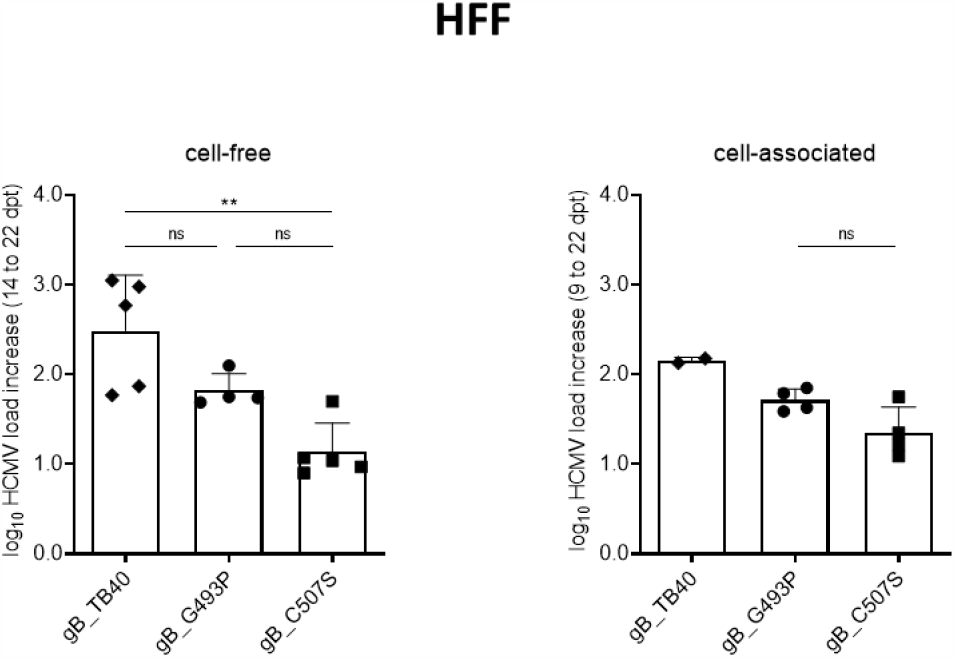
Early viral replication and release of viral DNA of gB mutants transfected into fibroblasts. The log_10_ increase in viral DNA load between 14 and 22 dpt and 9 and 22 dpt in cell-free and cell-associated samples, respectively, was calculated for all gB mutants. Mutants with the same gB mutation independent of the gO genotype (GT) background were grouped. Mean values with error bars indicate standard deviation (SD). **, p < 0.01; ns, not significant, one-way ANOVA with Tukey’s multiple comparisons test; HCMV, human cytomegalovirus; HFF, human foreskin fibroblast; dpt, days post transfection; GT, genotype.

**Supplementary Figure 3.**
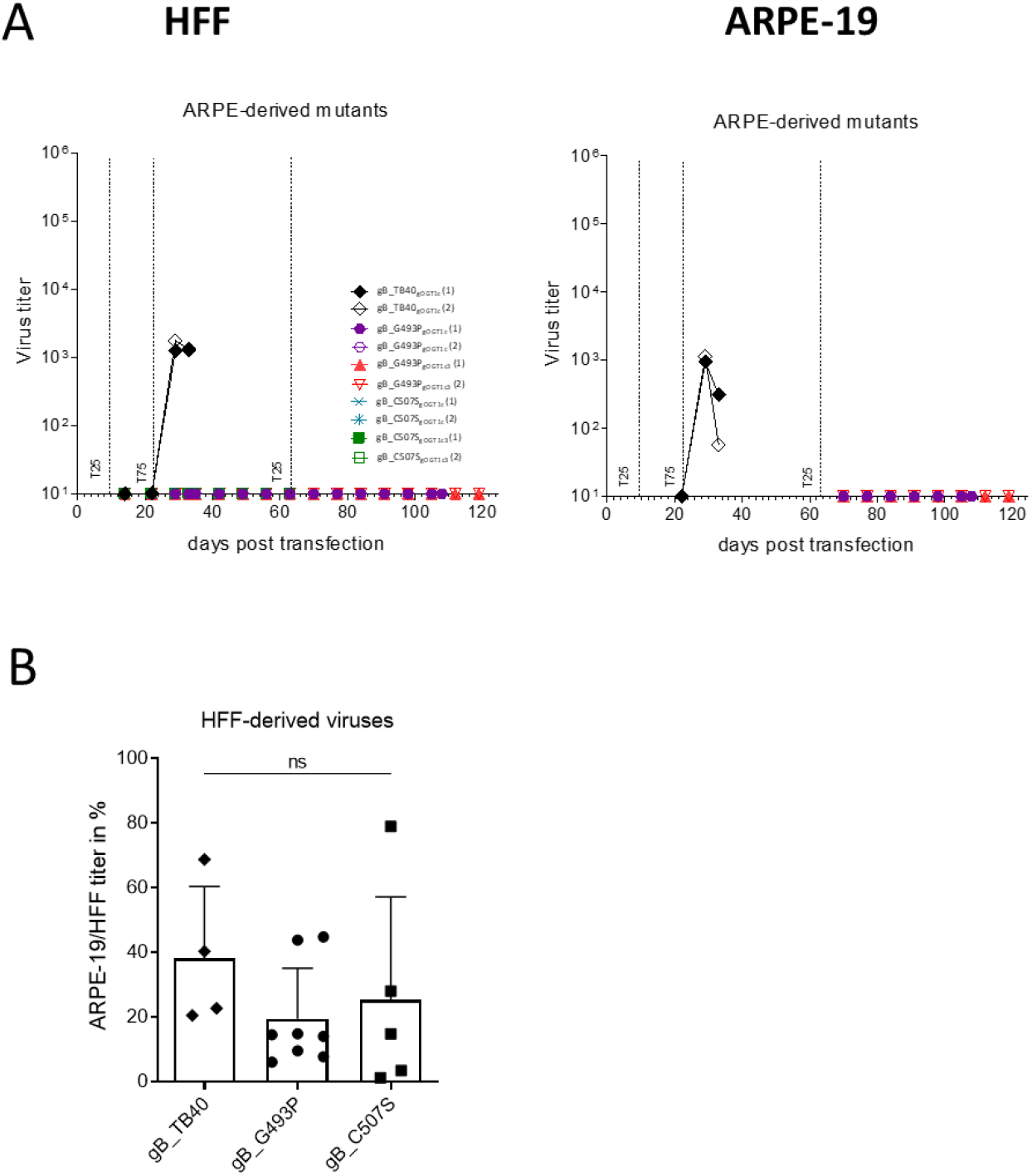
Infection efficiency and epithelial cell tropism. A) Epithelial cells (ARPE-19) were transfected with the respective TB40-BAC-luc DNA and long-term cultured. During reconstitution, cell-free samples were taken from the supernatant once a week during medium change. Subsequent to the harvest of all gB clones, infection efficiency of ARPE-derived viruses was tested in parallel through a luciferase assay. After infection, fibroblasts (HFF) were incubated for 48 h and ARPE-19 cells for 72 h. The virus titer (mean of duplicates) was determined through the conversion of the RLU to MOI and further into PFU. Parental gB_TB40gOGT1c in black, gB_G493P mutants in red and purple, and gB_C507S mutants in green and petrol, each in gOGT1c and gOGT1c3 background, respectively. ARPE-19-derived viruses of all cell-free samples were tested on HFFs (left). Positive samples on HFFs were tested on ARPE-19 cells (right). B) Relative ARPE-19 cell tropism of HFF-derived gB mutants. ARPE-19 to HFF infection ratio was calculated from all mutants which showed a detectable infectivity in both cell types (see Figure 2C). Mutants with the same gB mutation independent of the gO genotype (GT) background were grouped. Mean values with error bars indicate standard deviation (SD). Ns, not significant; one-way ANOVA. HCMV, human cytomegalovirus; HFF, human foreskin fibroblast; ARPE-19, retinal pigment epithelial cells; GT, genotype; T25, transfer into a T25 flask; T75, transfer into a T75 flask; RLU, relative light units; MOI, multiplicity of infection; PFU, plaque forming unit.

**Supplementary Figure 4.**
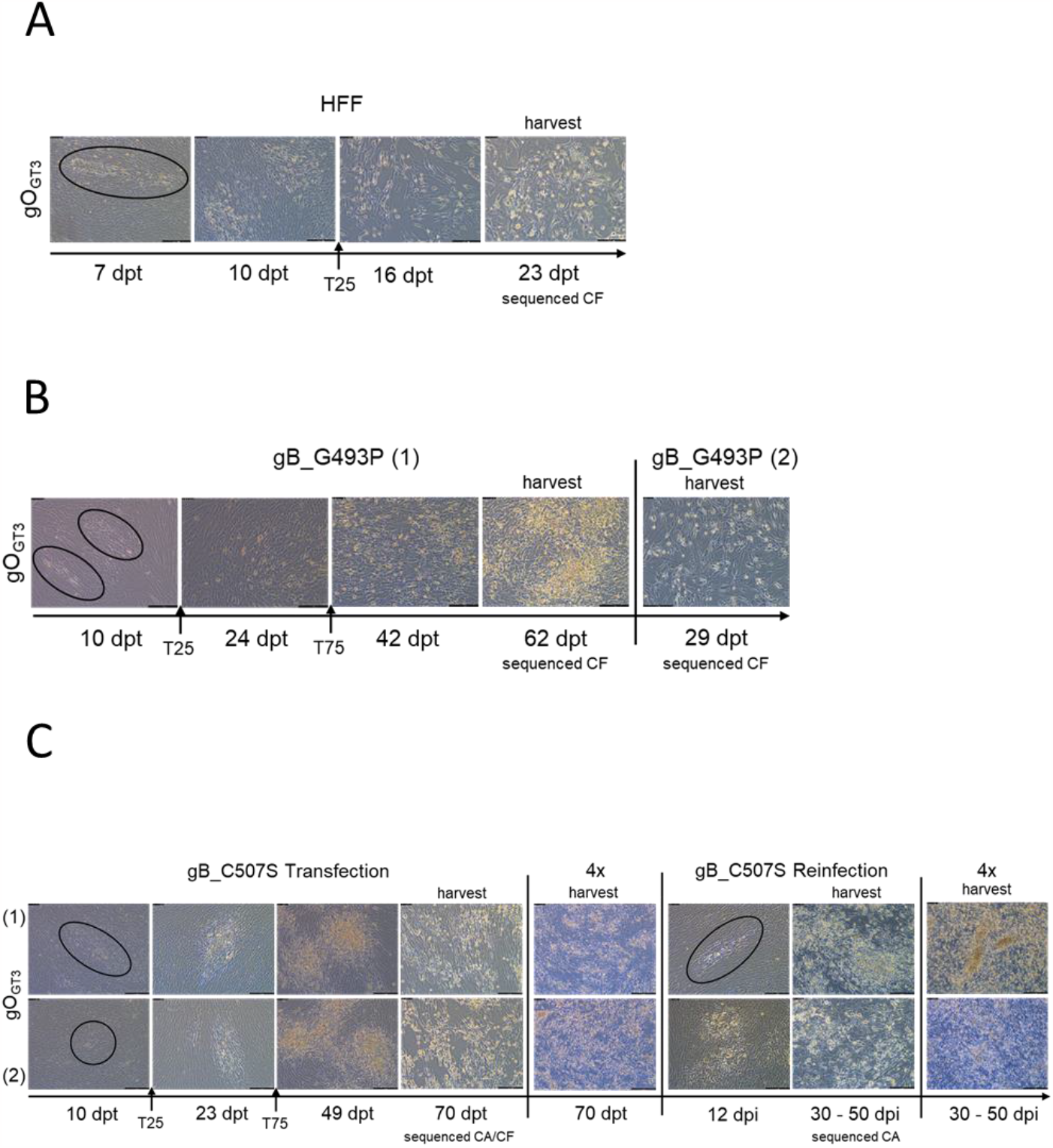
Spread morphology of gB_gOGT3 mutants. Human fibroblasts (HFF) were transfected with the respective TB40-BAC-luc DNA and long-term cultured. After transfection, pictures were taken every three to four days via light microscopy (Leica). Representative fields of each picture are shown at the indicated time points in 10X if not otherwise indicated. A) gB_TB40_gOGT3_ parental strain. Cell-free (CF) samples were sequenced at the time of harvest by whole genome sequencing (WGS). B) gB_G493P mutants. CF samples were WGS sequenced at the time of harvest. C) gB_C507S mutants. WGS of cell-associated (CA) and CF samples at time point of harvest with ∼ 80% cytopathic effect (CPE). HFFs were subsequently reinfected with gB_C507S CF samples from the time of harvest for further culturing (right) until 30 - 47 days post infection (dpi). CA samples were WGS sequenced at the time of harvest. HCMV, human cytomegalovirus; HFF, human foreskin fibroblast; T25, transfer into a T25 flask; T75, transfer into a T75 flask; (1), clone 1; (2), clone 2; dpt, days post transfection; dpi, days post infection; CF, cell-free; CA, cell-associated; CPE, cytopathic effect.

**Supplementary Table 1a.** Whole genome sequencing of cell-free (CF) and cell-associated (CA) fraction of BAC-derived parental strain and gB mutants at harvest time to confirm the presence of introduced mutation into gB domain (D) III α3 helix.

| # of gB mutants | gB mutant name | Clone | gO background | Cell type | SNP at nt position<sup>1</sup> | Introduced gB mutation | Mutation frequency % | Average genome coverage | Material | CPE | Harvest dpt / dpi* |
| --- | --- | --- | --- | --- | --- | --- | --- | --- | --- | --- | --- |
| P1 | gB_TB40<sub>gOGT1c</sub> | 1 | GT1c | HFF | — | — | — | 692 | CF | ~ 100% | 26 |
| P2 | gB_TB40<sub>gOGT1c</sub> | 2 | GT1c | HFF | — | — | — | 689 | CF | ~ 100% | 27 |
| P3 | gB_TB40<sub>gOGT1c</sub> | 3 | GT1c | HFF | — | — | — | 1229 | CA | ~ 100% | 22 |
| P4 | gB_TB40<sub>gOGT1c</sub> | 4 | GT1c | HFF | — | — | — | 1930 | CA | ~ 100% | 22 |
| P5 | gB_TB40<sub>gOGT1c3</sub> | 1 | GT1c3 | HFF | — | — | — | 341 | CF | ~ 100% | 23 |
| P6 | gB_TB40<sub>gOGT1c3</sub> | 2 | GT1c3 | HFF | — | — | — | 230 | CF | ~ 100% | 23 |
| P7 | gB_TB40<sub>gOGT3</sub> | 1 | GT3 | HFF | — | — | — | 409 | CF | ~ 100% | 23 |
| P8 | gB_TB40<sub>gOGT3</sub> | 2 | GT3 | HFF | — | — | — | 249 | CF | ~ 100% | 20 |
| P9 | gB_TB40<sub>gOGT1c</sub> | 1 | GT1c | ARPE-19 | — | — | — | 167 | CA | ~ 100% | 34 |
| P10 | gB_TB40<sub>gOGT1c</sub> | 2 | GT1c | ARPE-19 | — | — | — | 257 | CA | ~ 100% | 34 |
| 1 | gB_G493P<sub>gOGT1c</sub> | 1 | GT1c | HFF | 83588 | G493P | 100 | 586 | CA | ~ 100% | 29 |
| 2 | gB_G493P<sub>gOGT1c</sub> | 2 | GT1c | HFF | 83588 | G493P | 100 | 492 | CA | ~ 100% | 29 |
| 3 | gB_G493P<sub>gOGT1c3</sub> | 1 | GT1c3 | HFF | 83588 | G493P | 100 | 2178 | CA | ~ 100% | 29 |
| 4 | gB_G493P<sub>gOGT1c3</sub> | 2 | GT1c3 | HFF | 83588 | G493P | 100 | 631 | CA | ~ 100% | 29 |
| 5 | gB_G493P<sub>gOGT3</sub> | 1 | GT3 | HFF | 83588 | G493P | 100 | 723 | CF | ~ 100% | 62 |
| 6 | gB_G493P<sub>gOGT3</sub> | 2 | GT3 | HFF | 83588 | G493P | 100 | 527 | CF | ~ 100% | 29 |
| 7 | gB_C507S<sub>gOGT1c</sub> | 1 | GT1c | HFF | 83547 | C507S | 100 | 170 | CA | ~ 100% | 79 |
|  | gB_C507S<sub>gOGT1c</sub> | 1 | GT1c | HFF | 83547 | C507S | 100 | 1242 | CF | ~ 100% | 79 |
| 8 | gB_C507S<sub>gOGT1c</sub> | 2 | GT1c | HFF | 83547 | C507S | 100 | 543 | CA | ~ 100% | 93 |
|  | gB_C507S<sub>gOGT1c</sub> | 2 | GT1c | HFF | 83547 | C507S | 100 | 1185 | CF | ~ 100% | 93 |
| 9 | gB_C507S<sub>gOGT1c3</sub> | 1 | GT1c3 | HFF | 83547 | C507S | 100 | 603 | CA | ~ 100% | 114 |
|  | gB_C507S<sub>gOGT1c3</sub> | 1 | GT1c3 | HFF | 83547 | C507S | 100 | 1754 | CF | ~ 100% | 114 |
| 10 | gB_C507S<sub>gOGT1c3</sub> | 2 | GT1c3 | HFF | 83547 | C507S | 100 | 250 | CA | ~ 100% | 100 |
|  | gB_C507S<sub>gOGT1c3</sub> | 2 | GT1c3 | HFF | 83547 | C507S | 100 | 1241 | CF | ~ 100% | 100 |
| 11 | gB_C507S<sub>gOGT3</sub> | 1 | GT3 | HFF | 83547 | C507S | 100 | 423 | CA | ~ 80% | 70 |
|  | gB_C507S<sub>gOGT3</sub> | 1 | GT3 | HFF | 83547 | C507S | 100 | 1500 | CF | ~ 80% | 70 |
|  | gB_C507S<sub>gOGT3</sub> | 1 | GT3 | HFF | 83547 | C507S | 100 | 1046 | CA | ~ 100% | 30* |
| 12 | gB_C507S<sub>gOGT3</sub> | 2 | GT3 | HFF | 83547 | C507S | 100 | 255 | CA | ~ 80% | 70 |
|  | gB_C507S<sub>gOGT3</sub> | 2 | GT3 | HFF | 83547 | C507S | 100 | 458 | CF | ~ 80% | 70 |
|  | gB_C507S<sub>gOGT3</sub> | 2 | GT3 | HFF | 83547 | C507S | 100 | 358 | CA | ~ 100% | 50* |
| 13 | gB_G493P<sub>gOGT1c</sub> | 1 | GT1c | ARPE-19 | 83588 | G493P | 95 | 28 | CA | ~ 50% | 107 |
| 14 | gB_G493P<sub>gOGT1c3</sub> | 1 | GT1c3 | ARPE-19 | 83588 | G493P | 100 | 27 | CA | ~ 50% | 121 |
| 15 | gB_G493P<sub>gOGT1c3</sub> | 2 | GT1c3 | ARPE-19 | 83588 | G493P | 99 | 177 | CA | ~ 60% | 121 |

#, number; gB, glycoprotein B; gO, glycoprotein O; GT, genotype; SNP, single nucleotide polymorphism; nt, nucleotide; CA, cell-associated; CF, cell-free; D, domain; HFF, human foreskin fibroblast; ARPE-19, adult retinal pigment epithelial cells; CPE, cytopathic effect; dpt, days post transfection; dpi, days post infection. (1) numbering according to TB40-BAC4-luc sequence.

**Supplementary Table 1b.**
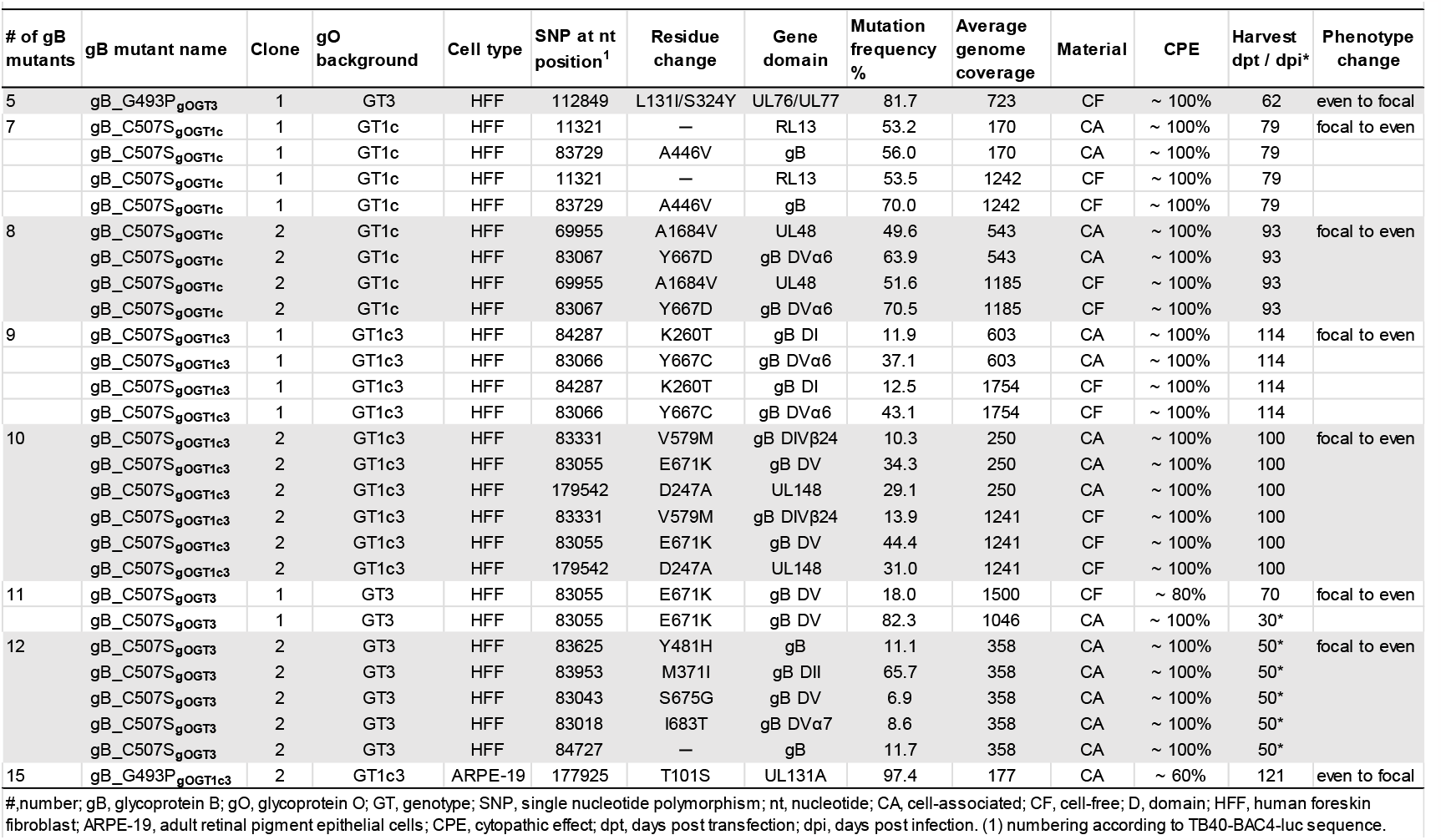
Whole genome sequencing of cell-free (CF) and cell-associated (CA) fraction of BAC-derived gB mutants at harvest time to investigate for the evolution of additional mutations along the human cytomegalovirus genome during long-term culturing.

**Supplementary Table 1c.** Amplicon sequencing of enriched gB region of cell-free (CF) fraction of gB_C507S_gOGT1c_ and gB_C507S_gOGT1c3_ mutants prior to (56 dpt) and shortly after the phenotype change (62 dpt) and at harvest time.

| # of gB mutants | gB mutant name | Clone | gO background | Cell type | SNP at nt position<sup>1</sup> | Residue change | Gene domain | Mutation frequency % | Average gB gene coverage | CPE | dpt |
| --- | --- | --- | --- | --- | --- | --- | --- | --- | --- | --- | --- |
| 10 | gB_C507S<sub>gOGT1c</sub> | 1 | GT1c | HFF | — | — | — | — | 4058 | ~ 50% | 58 |
|  | gB_C507S<sub>gOGT1c</sub> | 1 | GT1c | HFF | 83729 | A446V | gB | 8.7 | 3586 | ~ 50% | 62 |
|  | gB_C507S<sub>gOGT1c</sub> | 1 | GT1c | HFF | 83729 | A446V | gB | 67.7 | 9667 | ~ 100% | 79 |
| 11 | gB_C507S<sub>gOGT1c</sub> | 2 | GT1c | HFF | — | — | — | — | 3764 | ~ 50% | 58 |
|  | gB_C507S<sub>gOGT1c</sub> | 2 | GT1c | HFF | — | — | — | — | 6577 | ~ 50% | 62 |
|  | gB_C507S<sub>gOGT1c</sub> | 2 | GT1c | HFF | 83067 | Y667D | gB DVα6 | 70.66 | 706 | ~ 100% | 93 |
| 12 | gB_C507S<sub>gOGT1c3</sub> | 1 | GT1c3 | HFF | — | — | — | — | 5082 | ~ 50% | 58 |
|  | gB_C507S<sub>gOGT1c3</sub> | 1 | GT1c3 | HFF | — | — | — | — | 31241 | ~ 50% | 62 |
|  | gB_C507S<sub>gOGT1c3</sub> | 1 | GT1c3 | HFF | 84287 | K260T | gB DI | 17.0 | 4062 | ~ 100% | 114 |
|  | gB_C507S<sub>gOGT1c3</sub> | 1 | GT1c3 | HFF | 83066 | Y667C | gB DVα6 | 43.1 | 4062 | ~ 100% | 114 |
| 13 | gB_C507S<sub>gOGT1c3</sub> | 2 | GT1c3 | HFF | — | — | — | — | 13043 | ~ 50% | 58 |
|  | gB_C507S<sub>gOGT1c3</sub> | 2 | GT1c3 | HFF | — | — | — | — | 4124 | ~ 50% | 62 |
|  | gB_C507S<sub>gOGT1c3</sub> | 2 | GT1c3 | HFF | 83331 | V579M | gB DIVβ24 | 15.57 | 5239 | ~ 100% | 100 |
|  | gB_C507S<sub>gOGT1c3</sub> | 2 | GT1c3 | HFF | 83055 | E671K | gB DV | 41.49 | 5239 | ~ 100% | 100 |

#, number; gB, glycoprotein B; gO, glycoprotein O; GT, Genotype; D, domain; SNP, single nucleotide polymorphism; HFF, human foreskin fibroblast; CPE, cytopathic effect; dpt, days post transfection; (1) numbering according to TB40-BAC4-luc sequence.

**Supplementary Table 2.**
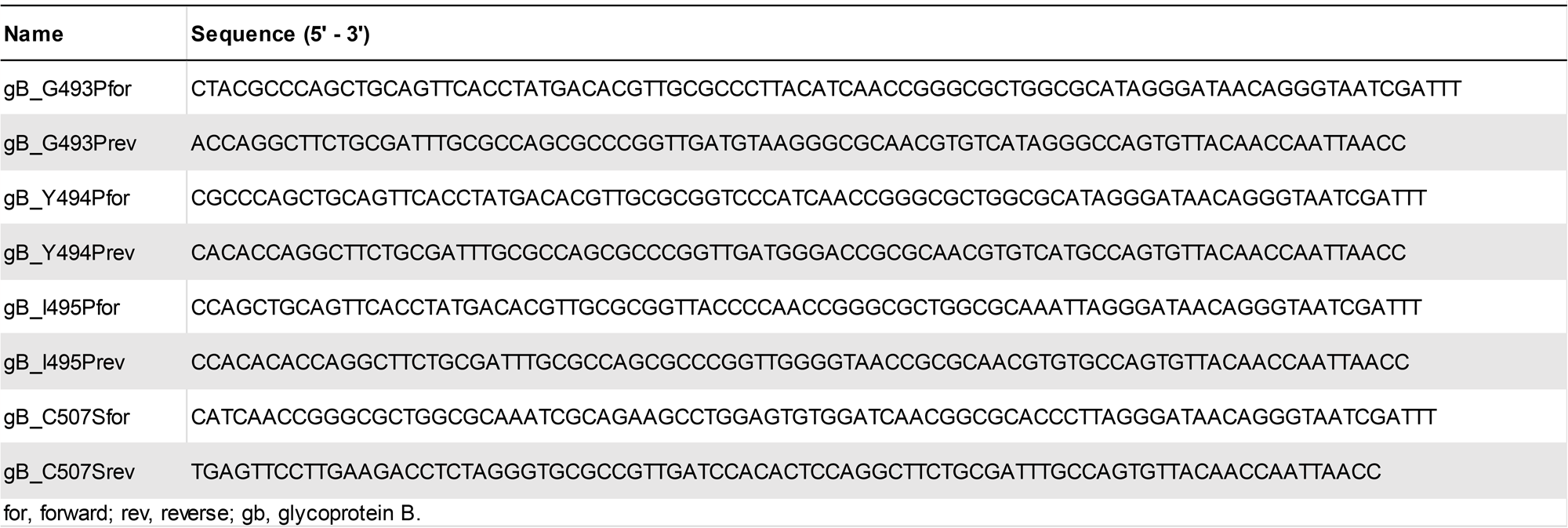
Primer for introduction of glycoprotein B (gB) point mutations by en passant mutagenesis.

